# Self-assembly vascularized human cardiac organoids model cardiac diseases in petri dishes and in mice

**DOI:** 10.1101/2023.08.26.554935

**Authors:** Qixing Zhong, Yao He, Li Teng, Yinqian Zhang, Ting Zhang, Yinbing Zhang, Qinxi Li, Bangcheng Zhao, Daojun Chen, Zhihui Zhong

## Abstract

In this study, we generated self-assembly cardiac organoids (COs) from human pluripotent stem cells by dual-phase modulation of Wnt/β-catenin pathway, utilizing CHIR99021 and IWR-1-endo. The resulting COs exhibited a diverse array of cardiac-specific cell lineages, cardiac cavity-like structures and demonstrated the capacity of spontaneous beating and vascularization *in vitro*. We further employed these complex and functional COs to replicate conditions akin to human myocardial infarction and SARS-CoV-2 induced fibrosis. These models accurately captured the pathological characteristics of these diseases, in both *in vitro* and *in vivo* settings. In addition, we transplanted the COs into NOD SCID mice and observed that they survived and exhibited ongoing expansion *in vivo.* Impressively, over a span of 75-day transplantation, these COs not only established blood vessel-like structures but also integrated with the host mice’s vascular system. It is noteworthy that these COs developed to a size of approximately 8 mm in diameter, slightly surpassing the dimensions of the mouse heart. This innovative research highlighted the potential of our COs as a promising avenue for cardiovascular research and therapeutic exploration.

## Introduction

Cardiovascular disease (CVD) ranks among the leading causes of death worldwide.(1, 2) Understanding the pathogenesis of CVD is crucial for both prevention and treatment strategies. It is thus imperative to study the pathophysiological mechanism underlying CVD and discover novel drugs for its treatment. A prerequisite to this is creating *in vitro* models that closely replicate the biological function of the human heart. These models will not only facilitate mechanistic studies on CVD but also open avenues for exploring the exogenous heart tissues or organ translational applications in CVD treatment.

In the past century, two-dimensional (2D, monolayer) cell lines and animal models have played crucial role in biomedical researches. However, these models have certain limitations: In the human body, cells live in a three-dimensional (3D) environment where they interact with neighboring cells and the extracellular matrix (ECM), (3, 4) while the 2D cell lines fail to replicate the physiological microenvironment or the intricate intercellular interactions found in the human body. As a result, they are unable to effectively simulate the complexity of human organs. (5, 6) Animal models have long been utilized in pharmaceutical researches; However, they come with their own drawbacks: the species disparity between animals and humans often results in the failure of translating promising pre-clinical findings to successful clinical trials. (3, 7)

Fortunately, the emergence of cutting-edge 3D-cell culture technology introduced a promising solution in the form of *in vitro* human organoids. With their 3D structure and intracellular microenvironment, human organoids closely mimic the structure and function of human organs. (8, 9) Consequently, human organoid models have emerged as optimal tools for studying disease mechanisms and conducting high-throughput drug screening in pre-clinical studies. Notably, the FDA has recently acknowledged the acceptance of results from organoid studies for pre-clinical studies preceding human trials. (10) This development presents a remarkable opportunity to leverage human organoids for more effective preclinical studies. As they can now serve as a “phase zero” of clinical trials, enabling the evaluation of drug candidates before phase 1 trials, thus reducing costs, time, and enhancing the overall efficiency of drug development. (11)

However, the utilization of *in vitro* human organoids is not without limitations. Unlike animal models, human organoids often lack functional vascular, nervous or immune system. (12) One potential solution is to transplant the *in vitro* human organoids into the animals to form a hybrid model. By integrating the transplanted human organoids into the animal’s blood circulation and metabolic system, they can better mimic the physiological environment of human body. Studies have demonstrated that *in vivo t*ransplanted organoids exhibit more mature ultrastructure than the *in vitro* cultured organoids. (13) Therefore, utilizing these transplanted models can provide a more clinically relevant representation of the drug treatment process within the human body, surpassing the drug screening capabilities of *in vitro* human organoids alone.

In summary, there is a growing interest in using *in vivo* transplanted human cardiac organoids (COs) for studying CVDs. In our study, we successfully derived human COs from pluripotent stem cells. They exhibited a wide range of cardiac-specific cell lineages and cavity-like structures, coupled with the capability for spontaneous beating and vascularization. Furthermore, we employed these COs to successfully simulate conditions such as human myocardial infarction and SARS-CoV-2 induced fibrosis, both *in vitro* and after transplantation into SCID mice. The transplantation of COs not only demonstrated their survival within the mice but also revealed the development of blood vessel-like structures that integrated with the host mice. Remarkably, the COs expanded to a size of approximately 8 mm in diameter, slightly exceeding the dimensions of the mouse heart. This groundbreaking research underscores the potential of COs as a promising avenue for advancing cardiovascular research and therapeutic exploration.

## Method

### Cell Culture

The human H9 embryonic stem cell line was obtained from Beina Biotechnology Co., Ltd. H9 cells was cultured with mTeSR^TM^1 (Stemcell Technologies, #85850, Canada) in a six-well plate coated with Matrigel (Corning, #354277, USA) at 37□ (1h incubation). After cell seeding, the H9 cells were cultured at 37□ with 5% CO_2_ and the culture medium was changed every 3-4 days (2 mL per well). When the cell confluence was above 70%, Accutase (Stemcell Technologies, #7920, Canada) was used to dissociated the cells for passaging. H9 cells were routinely (once a week) tested for the absence of mycoplasma contamination (Shanghai Life-iLab Biotech Inc, # AC16L061).

### Establishment of cardiac organoids *in vitro*

H9 cells were seeded at 3.5 × 104 cells/well in U-bottom ultra-low attachment 96-well plates (Qingdao AMA CO., LTD, #WP96-6CCUSH, China) with RPMI medium (Gibco^TM^, #21870075, USA) supplemented with B-27^TM^ (1:50 dilution, Gibco^TM^, #A1895601, USA), 6 μM CHIR99021 (Selleck, #S1263, USA), 7.5 μM Y-27632 (Sigma-Aldrich, #Y0503, USA) and 50 μg/mL Matrigel (Corning, #356231, USA). Cells were centrifuged at 1500 rpm/min for 10 min to force the 3-dimensional formation of the embryoid bodies (EBs). After 24 hours, EBs were cultured with RPMI+B27 medium (200 uL) and 1.5 μM CHIR99021 for 24 hours. After that, the EBs were cultured with RPMI+B27 medium for another 24 hours. On day 3, EBs were cultured with RPMI+B27 medium and 2.5 μM IWR-1-endo (Selleck, #S7086, USA) for 72 hours. Since day 6, the medium was refreshed with RPMI+B27 medium (200 uL per well) every 3-4 days.

### RT-qPCR analysis

Total RNA was extracted using trizol (Ambion, #15596026, USA). The cDNA was synthesized from 500 ng of total RNA using HiScript III RT SuperMix (Vazyme, #R323-01, China). RT-qPCR was performed using ChamQ Universal SYBR qPCR Master Mix (Vazyme, #Q711-02, China) on the CFX Connect Real-Time System (Bio-Rad, USA). The data were analyzed using the 2^^-ΔΔCt^ method. The list of the primer sequences used is provided in Supplementary Table 1.

**Supplementary Table 1.**
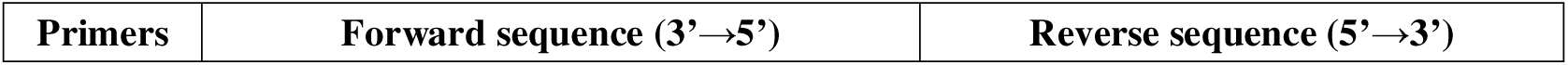
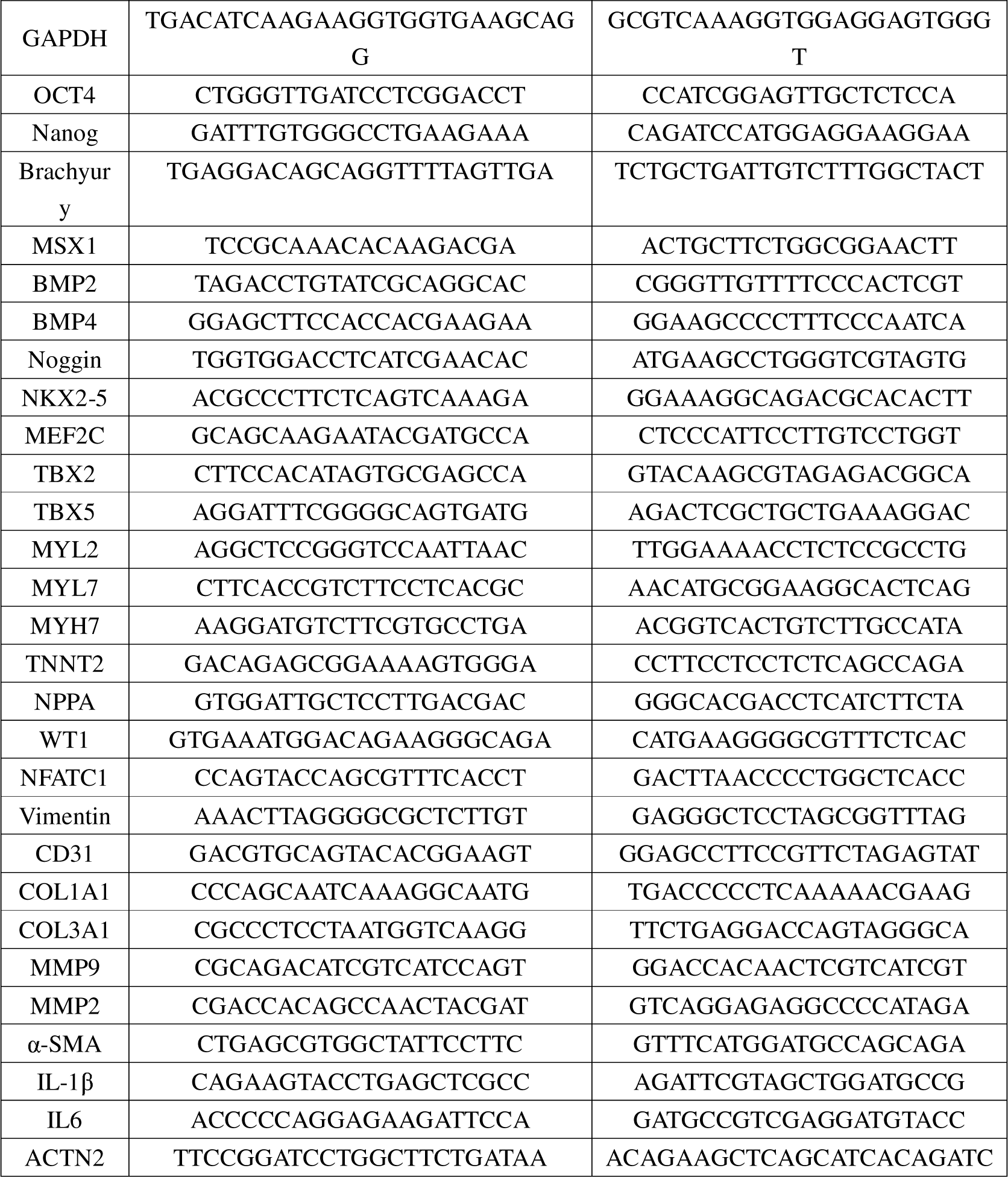
Primer sequences for RT-qPCR.

### Hematoxylin-eosin, TUNEL and immunofluorescence staining

COs were fixed with 4% paraformaldehyde (PFA, Thermo, #LEV138, USA) and embedded with paraffin. The embedded COs were sectioned into 2-4 μm thickness layers onto glass slides for Hematoxylin-eosin, TUNEL and immunofluorescence staining. The sections were dewaxed to water and antigenic repair was performed on the sections by thermal repair method. Goat serum was added to the sections and incubated at room temperature for 20 mins. Primary antibody (Supplementary Table 2) was then added to the sections and incubated overnight at 4□. On the next day, the sections were washed thrice with PBS (5 mins each). Subsequently, the secondary antibody (Supplementary Table 2) was added to the sections, incubated at 37□ for 30 mins, and then washed thrice with PBS (5 mins each). After that, DAPI (1:1000, Thermo, #D1306, USA) was added to the sections and incubated at room temperature for 10 mins. After staining, the sections were washed with PBS thrice (5 mins each). Finally, the sections were sealed with anti-fluorescence attenuation reagent (Solarbio, #S2100, China) and imaged using Inverted Confocal microscope (Andor dragonfly 200, UK). Images were analyzed with Imaris Viewer software (Oxford Instruments). Apoptosis rate from TUNEL staining was calculated as follows:

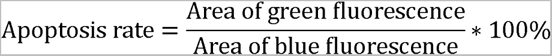

**Supplementary Table 2.** Antibodies used for immunofluorescence.

|  | Antibody name | Host Species | Dilution | Catalogue number | Manufacturer |
| --- | --- | --- | --- | --- | --- |
| Primary antibody | TNNT2 (cTnT) | Mouse | 1:100 | MA5-12960 | Invitrogen |
|  | NKX2-5 | Rabbit | 1:100 | 8792S | CST |
|  | MYL7 (MLC2a) | Mouse | 1:100 | 311011 | Synaptic Systems |
|  | MYL2 (MLC2v) | Rabbit | 1:100 | ab79935 | Abcam |
|  | WT1 | Rabbit | 1:100 | ab89901 | Abcam |
|  | Vimentin | Goat | 1:100 | ab11256 | Abcam |
|  | NFATC1 | Rabbit | 1:100 | ab25916 | Abcam |
|  | CD31 | Mouse | 1:100 | ab9498 | Abcam |
|  | CD31 | Rabbit | 1:100 | ab256569 | Abcam |
| Secondary antibody | FITC labeled (Green) | Goat anti-rabbit | 1:100 | GB22303 | Servicebio |
|  | CY3 labeled (Red) | Goat anti-mouse | 1:100 | GB21301 | Servicebio |

### Electrophysiological study of microelectrode array (MEA)

The cardiac organoids were transferred into a Cytoview MEA 24-well plate (Axion Biosystems, #m384-tmea-24w, USA) and placed in the center of the MEA to cover as many electrodes as possible. Only 20 to 50 μL medium was left in each well. The MEA plate was placed in the MEA instrument (Axion BioSystems, #MaestroEdge, USA) and equilibrated for 5 mins at 5% CO_2_ and 37□. The electrical signals of the COs were recorded for 2 mins at a time. Axis Navigator software version 3.7.3 was used for documentation and analysis.

### *In vitro* myocardial infarction model of cardiac organoids

COs were cultured with low glucose (1 g/L) RPMI+B27 medium and placed in a hypoxia incubator chamber (Billups-Rothenberg Inc., #MIC-101, USA) with an Oxygen analyzer (Nuvair O_2_ Quickstick, USA). Nitrogen gas was filled into the hypoxia incubator chamber to expel the oxygen until its level reached 1%. After that, the hypoxia incubator chamber was placed into incubator at 37□. The CellTiter-Glo^®^ 3D Cell Viability Assay Kit (Promega, #G9683, USA) was used to measure the Luminescence value of COs in each group. Micro-Image Analysis System software was used to record the beat rate of COs. The COs’ area were measured by the ImageJ (V1.8.0.112) software. The cell viability of COs from day 0 to day 5 after OGD modeling was calculated as follows:

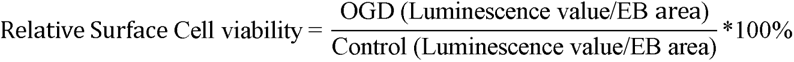

### *In vitro* myocardial fibrosis model of cardiac organoids

The spontaneous beating COs of day 20 were randomly divided into 6 groups: Control (without treatment), S protein (Bioss, #bs-46003P, China), S1 protein (Novoprotein, #DRA37, China), Ang □ (Aladdin, #A107852, China), S protein + Ang □ and S1 protein + Ang □ groups, with 10 COs in each group. The experimental period was 10 days and the medium was changed every three days. The beat count of each CO was measured daily and recorded. At the end of the experiment, COs were collected for further analysis.

### *In vivo* myocardial fibrosis model of cardiac organoids

NOD SCID mice (6-7 weeks old,18-20 g body weight, male and female) were purchased from Beijing Vital River Laboratory Animal Technology Co., Ltd. The animals were housed under specific pathogen-free (SPF) conditions, at 22 ± 2 °C and 55 ± 10% relative humidity, with a 12 h light/dark cycle. 10 COs (20 days) were subcutaneously transplanted into each mouse. After one week of transplantation, mice in each group were administered with SARS-CoV-2 spike protein (5 μg/mice/day, dissolved in sterile water), spike 1 protein (5 μg/mice/day), Ang II (1 μg/mice/day), spike protein + Ang II and spike 1 protein +Ang II, respectively, via intratracheal injection: mice were anesthetized via intraperitoneal injection with Pentobarbital Sodium (50 mg/kg) and were suspended vertically from their incisors. Gently pulled out the tongue with tweezers, pressed down on the tongue, inserted the endoscope probe into the mouth of mice, and observed its glottis, inserted the catheter into the trachea when the glottis opened, removed the catheter. After 10 days, the COs were extracted for further analysis. The above protocols used were approved by the Institutional Animal Care and Use Committee (IACUC) of West China Hospital, Sichuan University (Approval No. 20230301014). All animals were acclimated to the laboratory for one week.

### *In vivo* transplantation of cardiac organoids

NOD SCID mice (6-7 weeks old,18-20 g body weight, male and female) were purchased from Beijing Vital River Laboratory Animal Technology Co., Ltd. The animals were housed under specific pathogen-free (SPF) conditions, at 22 ± 2 °C and 55 ± 10% relative humidity, with a 12 h light/dark cycle. After *in vitro* culturing for 20 days, 20 COs were transferred from petri dish to the outer needle of the trocar. The trocar was punctured subcutaneously to the right back of mice and the COs were pushed out with the core of the needle. Mice were transplanted with blank COs (without DiR staining) or COs stained by 20 μM DiR (Beijing Fluorescence Biotechnology Co. Ltd, #22070, China; the screen of DiR concentration was shown in Supplementary Figure) for 30 mins. After transplantation, mice were firstly imaged by the optical In Vivo Imaging System (IVIS^®^ Lumia series III, PerkinElmer, USA). Subsequently, the mice were euthanized with 1.5% sodium pentobarbital (4 mL/kg). During extraction, the transplanted COs were isolated from the skin of the NOD SCID mice and placed into 24-well plates (Thermo, #142475, USA) with RMPI+B27 medium. The extracted COs were imaged with the IVIS, fixed with 4% PFA and analyzed by Hematoxylin-eosin staining and immunofluorescence analysis. The above protocols used were approved by the Institutional Animal Care and Use Committee (IACUC) of West China Hospital, Sichuan University (Approval No. 20230301014). All animals were acclimated to the laboratory for one week.

### Statistical analysis

GraphPad Prism software (GraphPad Software Inc., San Diego, USA) version 9.0 was used for statistical analysis. All data were expressed as mean ± standard deviation (SD) or standard error of the mean (SEM). The relative surface cell viability of COs was evaluated for statistical significance by the one-way analysis of variance (ANOVA) test. The beat rate of COs was evaluated by the two-way ANOVA test. Comparison of EB’s area among CO group and PT-CO group was tested by the Mean-Whitney test. *P* values < 0.05 were considered statistically significant.

## Results

### Establishment of self-assembly and functional COs from hPSCs *in vitro*

To create functional COs, we induced the cardiac differentiation of human pluripotent stem cells by two-phase modulation of Wnt/β-catenin pathway: At the embryoid body (EB) formation stage, this pathway was activated by the addition of CHIR99021 and sequential inhibited by the addition of IWR-1-endo for 3 days (Fig 1.A). (14) Spontaneously beat of the COs was observed from day 8 onwards and the COs were cultured for more than 80 days *in vitro*. (Fig 1.B). Their expansion reached an approximate area of 4×10^5^ μm^2^ by day 40, after which no substantial further growth was documented (Fig. 1C). This phenomenon could be due to maturing cardiac cells reducing their proliferation and prioritizing enhanced differentiation. (15, 16) Normally, they started to form heart-like chamber structures from day 20, as demonstrated by HE staining (Fig. 1D). The COs’ spontaneous beat rate varied between 45 to 132 beats per minute, with an average of around 85 beats per minute (Fig 1E), which is similar to the beat rate of human heart: between 60-100 beats per minute. (17) Electrophysiological assessment after EB dissociation confirmed cardiac subtypes with nodal, atrial and ventricular profiles of various quantities (Fig. 1F). To better monitor the differentiation process, we conducted analysis of gene expressions across distinct stages of differentiation. As the differentiation process unfolded, a noticeable decline in pluripotency markers (Nanog, OCT4) was observed (Fig 1 G). (18) In alignment with our expectations, mesodermal markers (Brachyury, MSX1, BMP2/4 and Noggin) exhibited a gradual decrease from day 20 to 80, (19–21) concomitant with a robust accumulation of cardiac progenitor cells markers such as NKX2-5 and MEF2C (Fig. 1H). (22) The above findings collectively demonstrated the successful establishment of self-assembly functional COs *in vitro*.

**Figure 1.**
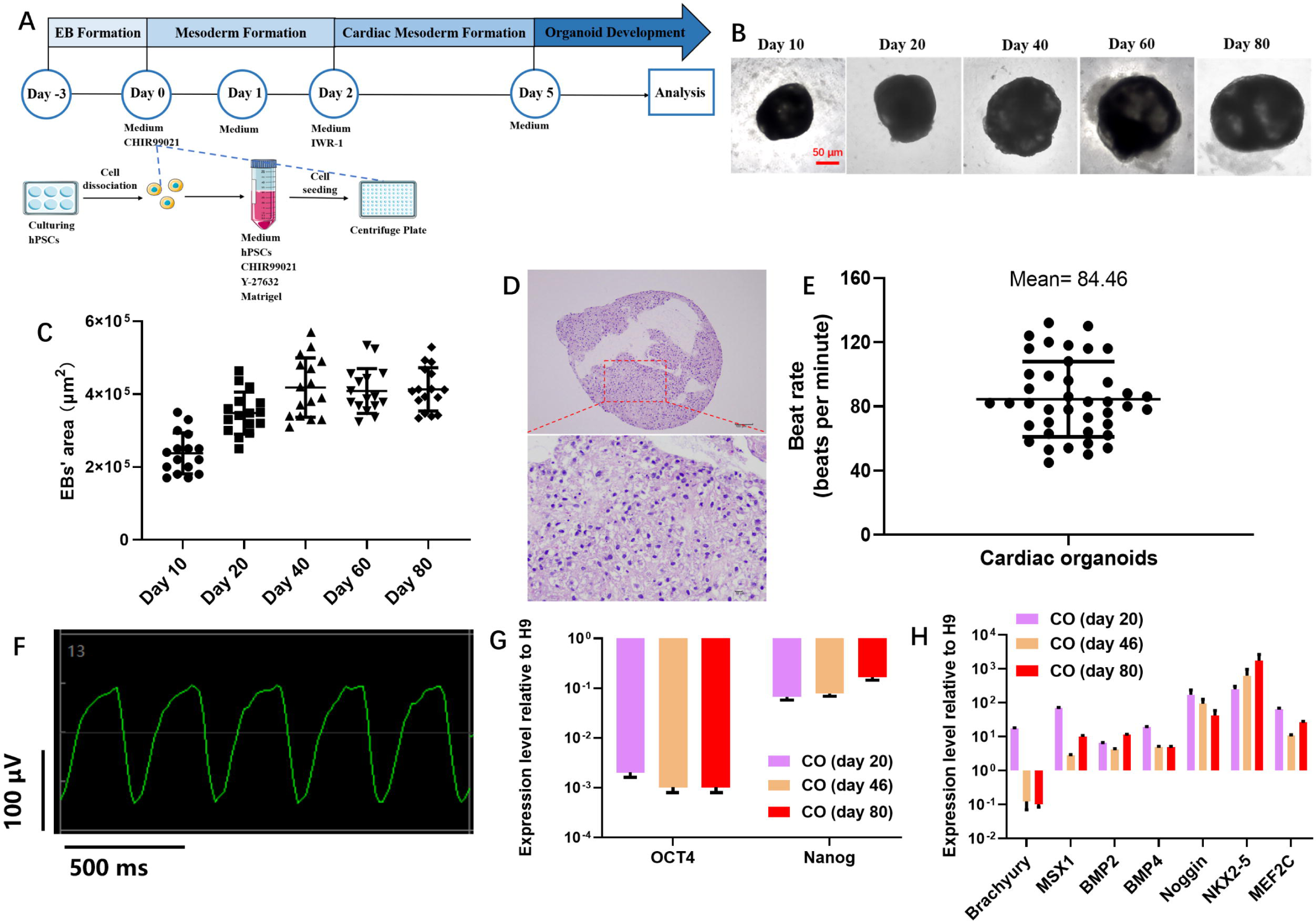
Establishment of self-assembly and functional COs from hPSCs *in vitro*. **(A)** Schematic diagram of COs derived from hPSCs. Representative bright field images (**B**) and size measurements (**C**) of COs at day 10, day 20, day 40, day 60 and day 80. n = 16. Scale bar: 50 μm. (**D**) Hematoxylin-Eosin staining images of CO at Day 20. Scale bar: 100 μm, inset: 10 μm. **(E)** Beat rate (per minute) of COs on day 25. n = 41. (**F**) Whole cell patch clamp recordings of multiple types of action potentials from CO derived cardiomyocytes. (**G**, **H**) RT-qPCR analysis of COs at day 20, day 46 and day 80. Gene expression level was normalized to the expression of GAPDH and H9. Data were presented as mean±SEM, n=3.

### COs generated atrial and ventricular specific cavities *in vitro*

We further analyzed cardiac-specific lineages of COs. Genes pivotal for cardiac specification, such as TBX2/5 and TNNT2, displayed a progressive increase in expression. (23–25) Moreover, there was a notable accumulation in the expression of MYL2 (MLC2v, encodes myosin light chain-2, ventricular subtype), MYL7 (MLC2a, encodes myosin light chain-7, atrial subtype), MYH7 and NPPA (Fig.2A). This suggests that our COs comprise cardiomyocytes that are committed to either atrial or ventricular lineages. (26–29) Subsequent immunofluorescence assessment verified the existence of NKX2-5, TNNT2, MYL2 and MYL7 expressing cells within the COs (Fig. 2B,C). Intriguingly, some MYL7-expressing cells were positioned around the chamber-like structure in the CO, suggesting the formation of an atrial-like structure within the myocardial cavities (Fig. 2D). (30)

**Figure 2.**
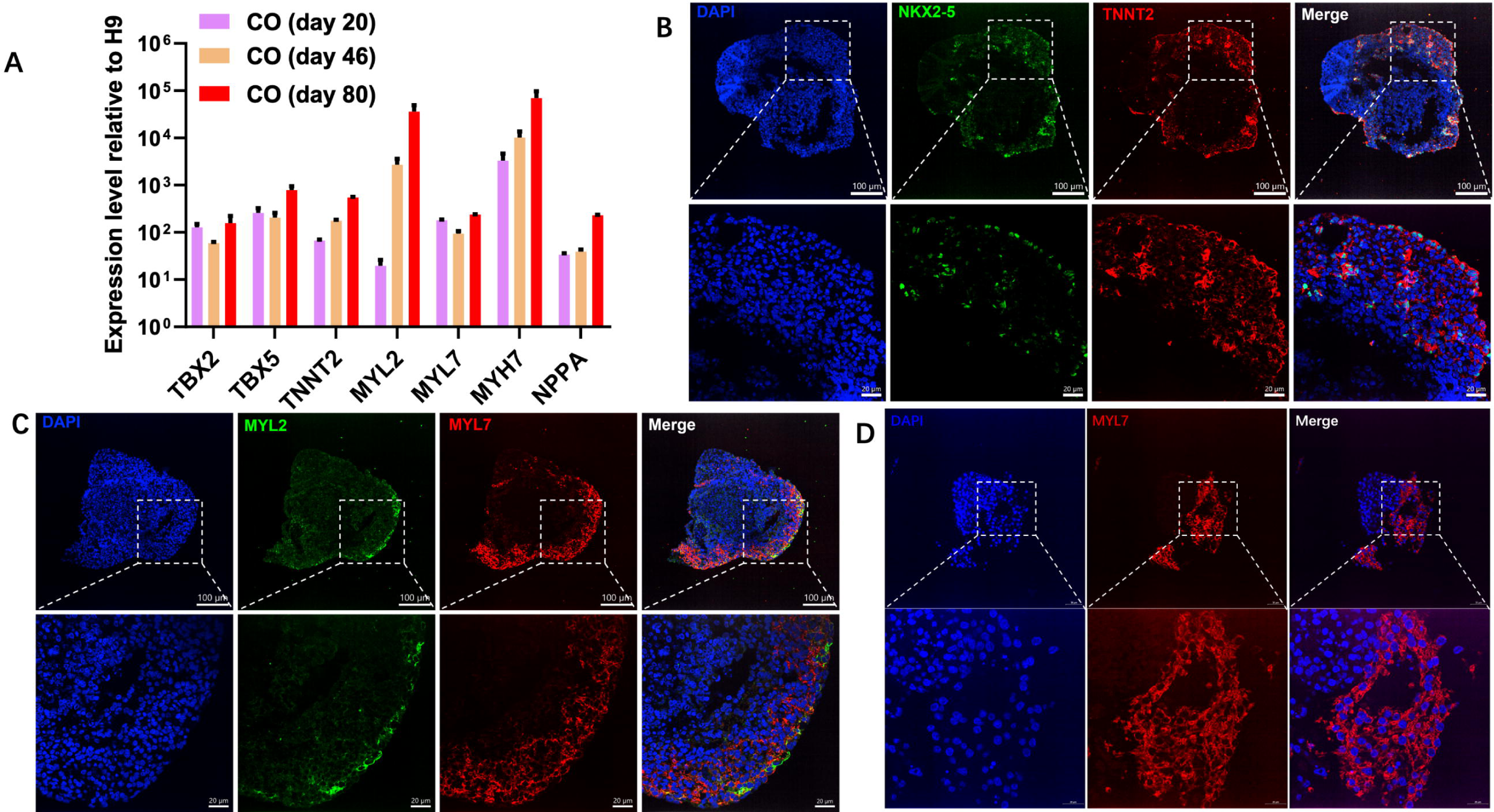
COs generated atrial and ventricular specific cavities *in vitro*. (**A**) RT-qPCR analysis of COs at day 20, day 46 and day 80. Gene expression level was normalized to the expression of GAPDH and H9. Data were presented as mean±SEM, n=3. Sectional immunofluorescence images of COs at day 30. (**B**) DAPI (blue), NKX2-5 (green), TNNT2 (red); (**C, D**) DAPI (blue), MYL2 (green), MYL7 (red); Scale bar: 100 µm or 50 μm, inset: 20 µm.

### COs generated multiple cardiac-specific structures and vascularized

Subsequent RT-qPCR analysis detected epicardial (WT1^+^), cardiac fibroblast (VIMENTIN^+^) and endocardium (NFATC1^+^) in our COs (Fig. 3A). (31, 32) Immunofluorescence identified Vimentin and TNNT2 positive cells in our COs, indicating the present of cardiac fibroblasts that also exist in human fetal heart (Fig. 3B). (33) Endocardial marker NFATC1 examination confirmed the formation of endocardial layer in our COs (Fig.3C). (35) Furthermore, confocal maximum intensity projection imaging highlighted the co-existence of epicardial (WT1) and cardiomyocytes (TNNT2) (Fig.3D). (36) Together, these findings demonstrated the structural integrity of our COs.

**Figure 3.**
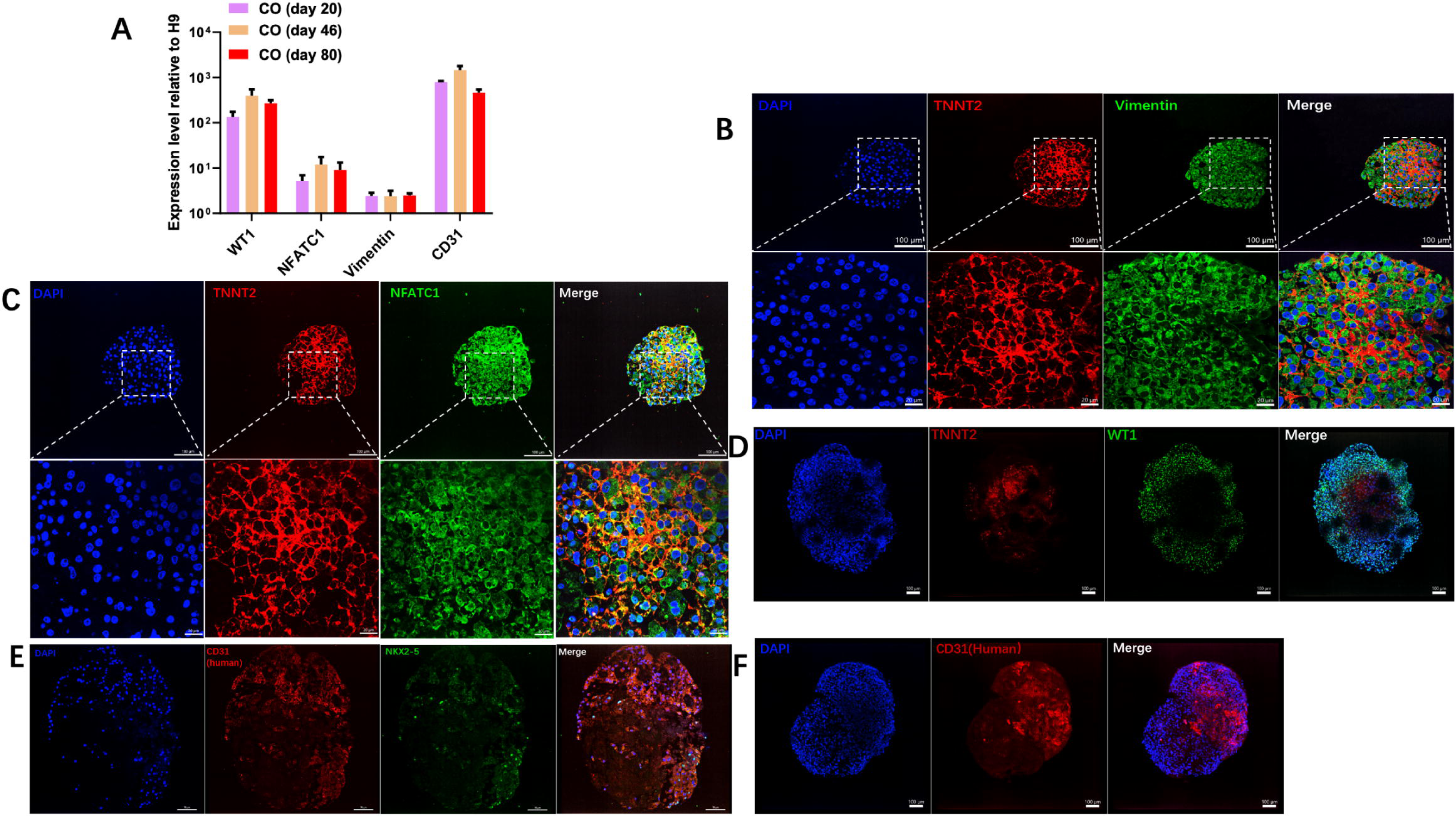
COs generated multiple cardiac-specific structures and vascularized. (**A**) RT-qPCR analysis of COs at day 20, day 46 and day 80. Gene expression level was normalized to the expression of GAPDH and H9. Data were presented as mean±SEM, n=3. Sectional immunofluorescence images of COs at day 30. (**B**) DAPI (blue), TNNT2 (red), Vimentin (green); (**C**) DAPI (blue), TNNT2 (red) and NFATC1 (green); (**E**) DAPI (blue), CD31(human, red) NXK2-5 (green). Scale bar: 100 µm or 70 µm, inset: 20 µm. Maximum intensity projection of COs immunofluorescence staining: (**D**) DAPI (blue), WT1 (green) and TNNT2 (red); (**F**) DAPI (blue), CD31(human, red). Scale bar: 100 µm.

Vascularization is crucial for human organoid development, ensuring adequate oxygen, nutrient, and metabolic compound exchange within an optimal diffusion range. (34) Notably, our RT-qPCR analysis showed increased expression of human endothelial cell marker CD31 (Fig. 3A). (35) Investigating further, both the sectional (Fig. 3E) and maximum intensity projection confocal images (Fig. 3F) confirmed the presence of the human CD31 positive cells, suggesting spontaneous vascularization within the COs.

### COs modelled myocardial infarction and SARS-CoV-2 induced fibrosis

To showcase the potential of COs in replicating human heart cardiac pathologies, we utilized them to simulate the condition of myocardial infarction and the damage caused by SARS-Cov-2 spike protein during COVID-19 infection. During heart attack, blocked artery limits oxygenated blood flow to the downstream myocardium, leading to significant cell death and reducing the heart’s ability to pump blood efficiently.(36) To model this *in vitro*, the COs of day 40 were cultured at 1% of O_2_ and 1g/L of glucose supply for 5 days. Their beat rate decreased, stopping entirely by day 5 in the oxygen-glucose deprivation (OGD) group (Fig. 4A). The surface cell viability declined accordingly, indicating a cell death in the exterior of the COs (Fig. 4B). During heart attack, the human body initiates compensatory efforts through the nervous system to restore cardiac output, primarily via adrenergic stimulation by norepinephrine (NE). (36) To mimic this, we treated our OGD model with 1μM of NE (from day 0) and the results showed the beat rate increased in comparison to the OGD group, which was reversed when cultured with 10 μM metoprolol, a beta-adrenergic blocker, (37) added from day 0 or on day 5, respectively (Fig. 4C). Furthermore, both the OGD and the OGD+NE organoids showed apoptotic TNUEL+ staining predominantly at the center of organoid sections, attributed mainly to the compromised oxygen level at the core of the COs. Intriguingly, we observed that the OGD+NE showed higher degree of apoptotic death than OGD group (Fig. 4D), as NE alone also causes apoptotic cell death. (38)

**Figure 4.**
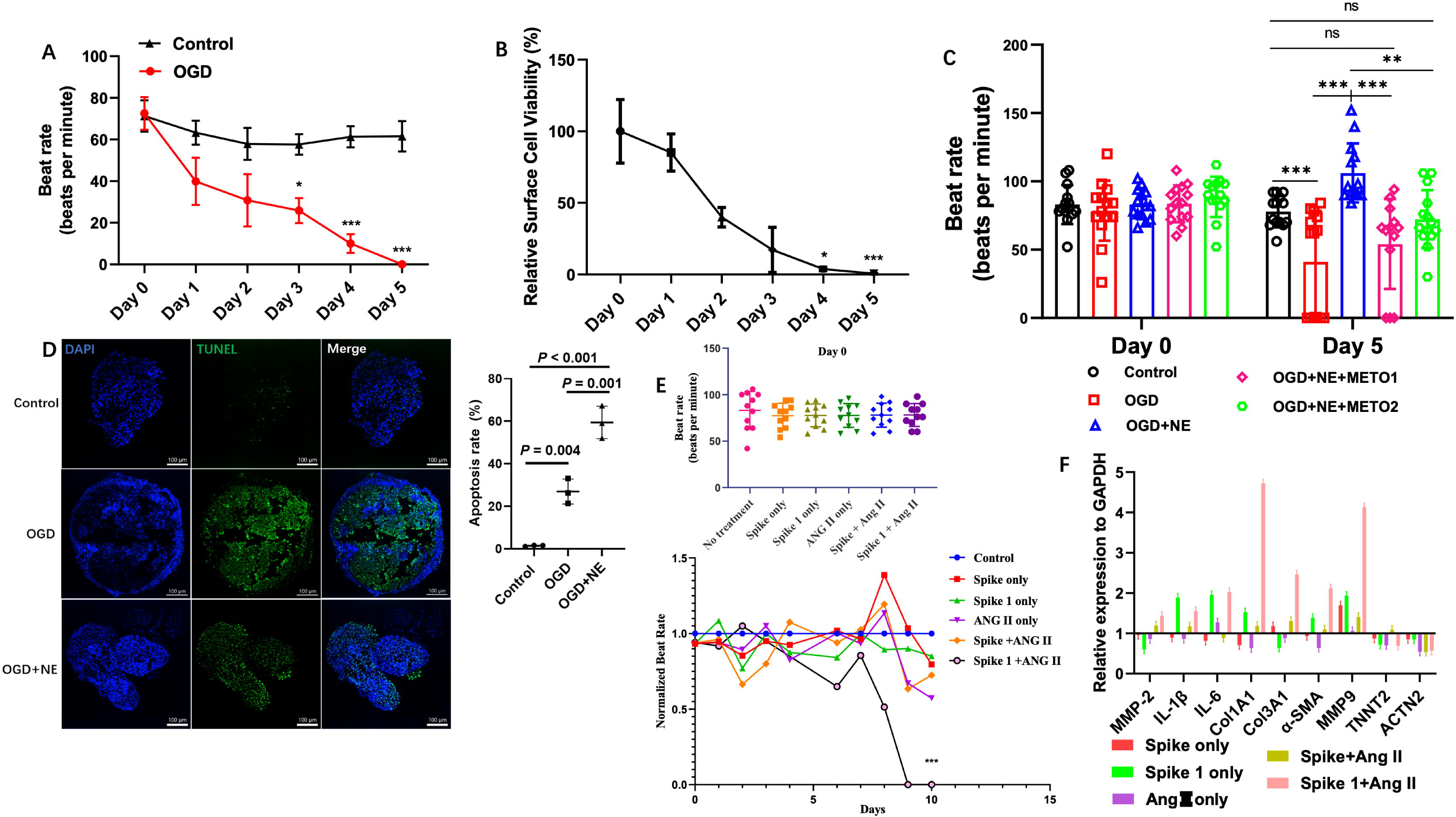
COs modelled myocardial infarction and SARS-CoV-2 induced fibrosis. (**A**) Beat rate (per minute) of COs from day 0 to day 5. Data were presented as mean±SEM, n = 4. (B) Relative surface cell viability to control of COs from day 0 to day 5 after OGD modeling. Data were presented as mean±SEM, n = 4. ***:p<0.001. (**C**) Beat rate (per minute) of COs in control, OGD, OGD+NE and OGD+NE+METO groups (METO1: added from day 0-5, METO2: added on day 5). (**D**) TUNEL apoptosis staining and TUNEL index quantification of COs in the control, OGD and OGD+NE groups on day 5. DAPI (blue) and TUNEL (green). Scale bar: 100 µm. (**E**) Beat rate count on day 0 and the normalized (to control) beat rate from day 0-10 of control, Spike protein (10 μg/mL), Spike 1 protein (10 μg/mL), Ang II (1μg/mL), Spike + Ang II and Spike 1+ Ang II groups. n=3, ***:p<0.001 to control. (**F**) RT-qPCR analysis of post transplantation COs, gene expression levels were normalized to the expression of GAPDH and control. Data were presented as mean±SD, n=3,

Long-COVID, lasting beyond four weeks after SARS-CoV-2 infection, significantly impacts individuals’ lives in the aftermath of COVID-19. It leads to notable cardiovascular issues, including dyspnea, palpitations, chest pain and arrhythmias. (39, 40) Among these issues, the SARS-CoV-2 spike protein causes cardiac fibrosis and reduced myocardial contractile in obese mice and in hiPSC-derived cardiomyocytes. (41, 42) Subsequently, we exposed the COs to spike proteins from Omicron and Wuhan variants, along with Angiotensin II (Ang II), which affects blood pressure and aldosterone release. (43) From the results, decline beat rate was observed after 7-day of treatment and in the Spike protein + Ang II group, the beat rate stopped entirely on day 9 (Fig. 4E). Subsequently, we transplanted the COs into NOD SCID mice and administrated the mice with similar treatments 10 days. RT-qPCR analysis of the post-transplanted COs confirmed the loss of cardiac muscle cells, evidenced by lower expression of TNNT2 and ACTN2. (41) Meanwhile, the fibrosis-related genes such as COL1A1, COL3A1, MMP-9, MMP-2 and α-SMA, (44–46) and genes that are related to inflammation: IL-1 β and IL-6 were detected to be upregulated after the spike 1+ang II treatment (Fig. 4F), which aligned with the physiological response within the human body. (47) Above results collectively demonstrated that COs have the capability to mimic human cardiac diseases and reproduce their pathological states in both *in vitro* and *in vivo* settings.

### COs developed in SCID mice and integrated with the host’s vasculature

To assess if COs remain alive and functional in SCID mice without culture medium, we transplanted them subcutaneously. COs were pre-stained with DiR to aid the localization. Fluorescence was later detected in the transplanted mice (Fig. 5A). After 14 days of transplantation, the COs were extracted and still showed detectable fluorescence (Fig. 5B). Unexpectedly, the post-transplantation CO (PT-COs) expanded to over 1 mm in diameter, three times larger than the simultaneously *in vitro* cultured control (Fig. 5C). The sectional HE staining of the PT-COs exhibited the cardiac muscle structures as well as blood vessels structure (Fig. 5D). Further sectional immunofluorescence analysis confirmed the PT-COs with MYL7 expression (Fig. 5E) and there were both mouse blood vessels (Fig. 5F) and human blood vessels (Fig. 5G) in them. Remarkably, after 75 days of transplantation, PT-COs grew to around 8 mm in diameter, 164 times larger than the *in vitro* cultured control. Furthermore, blood vessel-like structure on the PT-COs were observed (Fig. 5H). To determine the origin of these blood vessels, we performed maximum intensity projection immunofluorescence and human vessel-like tube formation across the organoid’s epicardial layer was detected (Fig. 5I). Taken together, these findings provide compelling evidence that the *in vivo* transplantation of the COs effectively facilitated their vascularization, leading to sustained development within the host mice.

**Figure 5.**
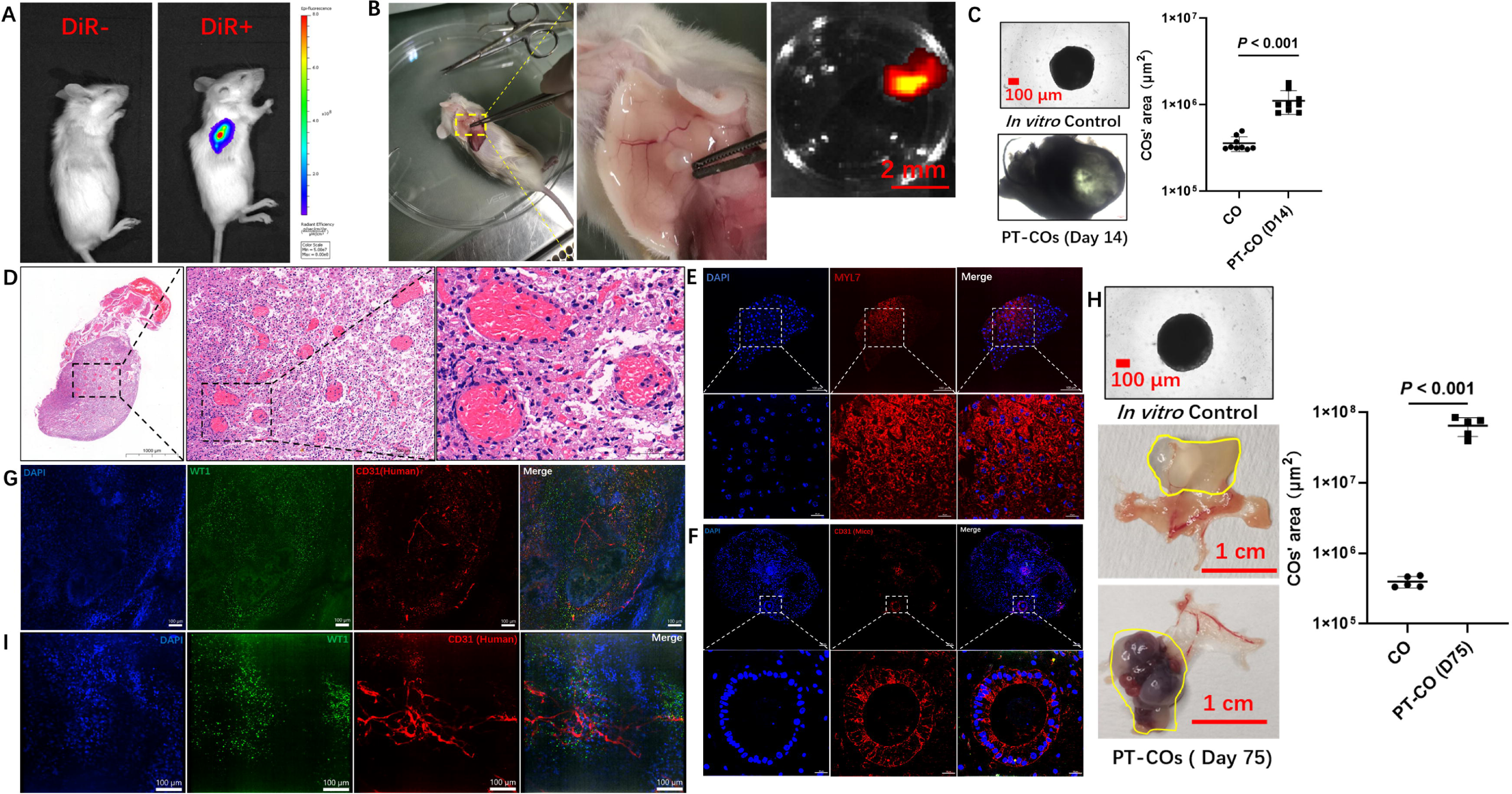
COs developed in SCID mice and integrated with the host’s vasculature. (**A**) IVIS images of SCID mice with COs (-/+ DiR staining) transplantation. (**B**) After subcutaneous transplantation (14 days or 75 days), COs with fluorescence were extracted from SCID mice. (**C**) The representative BF images of *in vitro* cultured COs and PT-COs of day 14. Scale bar: 100 µm. The area of the COs were measured using ImageJ, n= 9. (**D**) Hematoxylin-Eosin staining images of CO extracted from SCID mice. Scale bar: 1000 μm, inset: 200 μm and 50 µm. (**E**, **F**, **G**) Sectional immunofluorescence images of PT-COs of day 14. DAPI (blue), MYL7 (red), CD31(mice, red), CD31(human, red) and WT1 (green). Scale bar: 100 µm, inset: 20 µm. (**H**) The representative BF images of *in vitro* cultured COs and PT-COs of day 75. Scale bar: 100 µm. The area of the COs were measured using ImageJ, n=5. (**I**) Maximum intensity projection immunofluorescence staining of COs, WT1 (green), CD31 (human, red). Scale bar: 100 µm.

## Discussion

In recent years, hPSC-derived cardiomyocytes have become critically useful tools to model aspects of heart development, human genetic cardiac disease, therapeutic screening and cardiotoxicity testing. (35) Nonetheless, the complex structural morphology, multitude of tissue types present in the human heart and lack of vascularization impose severe limitations on current *in vitro* models. Current vascularization of cardiac organoids involves co-culture with rat endothelial cells or with vascular organoids.(48, 49) In our study we were able to establish the functional and spontaneously vascularized COs with cardiac chamber-like structures by regulating the Wnt/β-catenin pathway, with only two small molecules: CHIR99021 and IWR-1-endo. This method simplified the current protocols and enabled the establishment of complex, functional and vascularized COs on a larger scale *in vitro*.

SARS-CoV-2 Spike protein causes cardiac fibrosis, reduces myocardial contractile in obsess mice and causes damages in hiPSCs-derived cardiomyocytes. (41, 42) Our work, for the first time, has replicated the effects of the spike protein on the human heart using COs, both *in vitro* and after transplanted into mice. Due to the possessing of diverse cardiac cell types, intricate intercellular microenvironment and the presence of cardiac chamber-like structures in the COs, our findings hold greater clinical relevance than the animal or cell-only studies. This enables us to better understand the mechanisms behind spike protein-induced cardiac fibrosis and explore potential therapeutic approaches in the human context.

. Our transplantation of COs into NOD SCID mice discovered that, after subcutaneous transplantation, the COs survived in the mice by integrating with the host’s vascular system and formed vessel-like structure inside and around the COs. Although Lee et al. (2022) have reported similar subcutaneous transplantation of COs, (50) our study revealed that after longer time of transplantation (up to 75 days), the PT-COs expanded significantly (Fig. 5H), with dimensions marginally surpassing the normal mouse heart, which typically measures around 6 mm in diameter. (51) We hypothesize that the mouse’s circulatory system delivered essential nutrients that facilitated the growth, expansion and vascular maturation of the COs *in vivo*. In contrast, the COs cultured simultaneously in petri dish maintained significantly smaller size (Fig. 5H). This result reinforces the notion that the animal body, often dubbed ’the ultimate petri dish’, offers an ideal setting for organoid growth, encouraging further xenograft research with diverse human organoids. To delve deeper, future investigations such as RNA-seq analysis will be necessary to scrutinize the differences between *in vitro* and *in vivo* cultured COs. This analysis aims to identify pivotal signaling pathways critical to this observed phenotype.

The successful engraftment and integration of COs in mice offers numerous potential applications. For instance, we could establish various cardiac diseases models *in vitro* (beyond OGD or spike protein-induced damage) and then transplant them into animals. These *in vivo* COs models can be used for drug trials, applying different delivery methods like oral or intravenous administration in the animals. This strategy offers promising avenue for preliminary *in vivo* pharmacokinetic studies of drugs using human organoids, which is more clinically relevant than studies solely conducted on animals or *in vitro* human organoid. Additionally, these models could be employed for drug toxicity assessment, examining not just drug effects but also impacts of their metabolites.

Last but not least, achieving human COs comparable in size to a mouse heart that’s vascularized and operational within the mouse offers a chance to delve into its therapeutic promise. For example, these COs might be utilized to boost blood circulation. We envision creating *in vitro* COs from patient-specific induced pluripotent stem cells (iPSCs) and then transplanting them back into the same patient for novel therapeutic approaches. This pioneering method could present significant therapeutic solutions, potentially revolutionizing heart failure treatments.

## Author contributions

ZZ designed the entire study and supervised all segments of the study. QZ helped to design the research and prepared the manuscript. QZ and YH conducted all the experiments. LT, TZ, YZ helped to conduct the experimental procedures. YZ, QL, BZ and DC conducted the statistical analysis and revised the manuscript. The final manuscript was carefully reviewed and approved by all authors.

## Acknowledgement

The authors would like to thank Sichuan Junhui Biotechnology Co., Ltd. for providing the experimental platform for this research.

