## Supplementary figures and images for "Self-assembly vascularized human cardiac organoids model cardiac diseases in petri dishes and in mice"

### NA

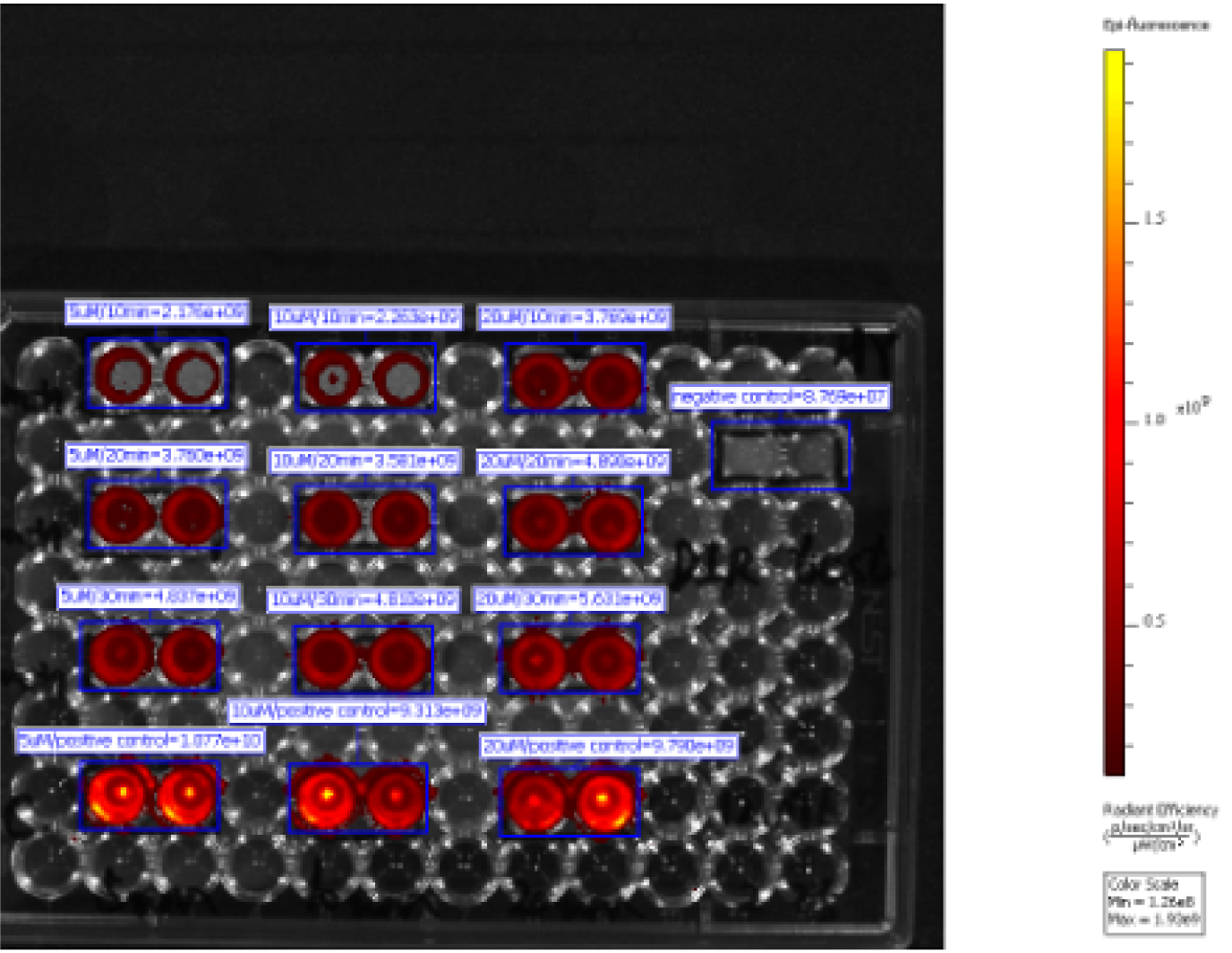
